# Tau protein in smooth muscle cells and tissues

**DOI:** 10.1101/2020.10.15.341867

**Authors:** Nataliia V. Shults, Sarah Seeherman, Nurefsan E. Sariipek, Vladyslava Rybka, Lucia Marcocci, Sergiy G. Gychka, Yasmine F. Ibrahim, Yuichiro J. Suzuki

## Abstract

Tau is a microtubule-associated protein and plays a critical role in the pathophysiology of neurons. However, whether tau protein is expressed in smooth muscle cells is unknown. Here, we report that tau protein is expressed and is constitutively phosphorylated at threonine 181 in various smooth muscle cell types, including human cerebral artery smooth muscle cells, human pulmonary artery smooth muscle cells, and human bronchial airway smooth muscle cells. We also detected the expression of tau protein in the vascular smooth muscle of brain tissues from patients with systemic hypertension who died of ischemic stroke. Immunofluorescence staining revealed that phosphorylated tau at threonine 181 is more organized in the cell than does total tau protein. A protein phosphatase inhibitor, calyculin A induced the formation of higher molecular weight species of phosphorylated tau as visualized by Western blotting, indicating the occurrence of tau aggregation. Immunofluorescence also showed that calyculin A caused the aggregation of phosphorylated tau and disrupted the cytoskeletal organization. These results demonstrate the existence of tau protein in smooth muscle cells and tissues and that smooth muscle tau is susceptible to protein phosphorylation and aggregation.

## Introduction

Tau is a microtubule-associated protein and a natively unfolded protein lacking a significant amount of secondary structure (Grundke-Iqbal et al., 1986). Tau promotes the self-assembly of tubulin into the microtubule and plays an important role in microtubule stabilization in the cell (Binder et al., 1985; Qiang et al., 2018). In physiological conditions, tau as a stabilizer of microtubules regulates cell differentiation and proliferation (Evans et al., 2018; Drubin & Kirschner, 1986; Weingarten et al., 1975). However, under pathological conditions, tau proteins assembly into insoluble aggregates, a hallmark of most neurodegenerative diseases referred to as tauopathies (Saha et al., 2019; Mudher et al., 2017; Mukrash et al., 2017; Narasimhan et al., 2017). The abnormal deposition of misprocessed and aggregated tau proteins in the neuromuscular system contributes to the development of supranuclear palsy, corticobasal degeneration, Pick’s disease, Huntington’s disease, argyrophilic grain disease, frontotemporal dementia and parkinsonism linked to chromosome 17, and globular glial tauopathy (Vuono, et al., 2015; Fernández-Nogales et al., 2018; Ballatore et al., 2007; Chin & Goldman, 1996; Feany & Dickson, 1996; Komori, 1999; Poorkaj et al., 1998). Also, the tau protein aggregation in the brain intracellular fibrils (neurofibrillary tangles) is a major hallmark of Alzheimer’s disease, the most common neurodegenerative dementia (Rossi et al., 2013; Rossi et al., 2008; Ittner et al., 2010; Johnson & Hartigan, 1999; Braak & Braak, 1991). Phosphorylation of tau contributes to disease-associated tau toxicity (Wang & Mandelkow, 2016; Šimić et al., 2016; Ittner et al., 2010; Khurana et al., 2006; Steinhilb et al., 2007). Tau is also expressed in non-neural cells including fibroblasts and lymphocytes (Ingelson et al., 1996; Thurston et al., 1996).

In many neurodegenerative disorders, the vascular component has a significant impact on brain metabolism and homeostasis. The loss of vasculature smooth muscle cells in Alzheimer’s disease impairs the clearance of β-amyloid and other toxic metabolites (ElAli et al., 2013; Hawkes et al., 2013; Meyer et al., 2008; Ervin et al., 2004).

Since it is not known if tau protein is expressed in smooth muscle cells and tissues, the present study examined the expression of tau protein in the smooth muscle. We found that various types of smooth muscle cells express tau protein that is subjected to protein phosphorylation and aggregation.

## Materials and Methods

### Histological measurements of human brain

Postmortem brain tissues were collected from patients with a history of systemic hypertension and who died of ischemic stroke. The tissues were taken from the perifocal zone of the ischemic infarct in region of the middle cerebral artery of the frontal lobe. Clinical studies were approved by the regional committee for medical research ethics in Kiev, Ukraine (ethical code: 81, 2016) and performed under the Helsinki Declaration of 1975 revised in 2013 or comparable ethical standards.

Brain tissues were immersed in buffered 10% formalin at room temperature and embedded in paraffin. Paraffin-embedded tissues were cut and mounted on glass slides. Tissue sections were subjected to immunohistochemistry using the Tau antibody (MilliporeSigma, Burlington, MA, USA).

### Cell Culture

Human brain vascular smooth muscle cells, human pulmonary artery smooth muscle cells, and human bronchial smooth muscle cells were purchased from ScienCell Research Laboratories (Carlsbad, CA, USA). Cells were cultured in accordance with the manufacturer’s instructions in 5% CO_2_ at 37°C. Cells were treated with calyculin A or H_2_O_2_ purchased from MilliporeSigma. For siRNA experiments, cells were transfected with an siRNA Transfection Reagent and control, Tau (h) or Tau (h2) siRNA from Santa Cruz Biotechnology (Dallas, TX, USA). 48 hours later, cell lysates were prepared.

### Western blotting

To prepare cell lysates, cells were washed in phosphate buffered saline and solubilized with lysis buffer containing 50 mM Hepes (pH 7.4), 1% (v/v) Triton X-100, 4 mM EDTA, 1 mM sodium fluoride, 0.1 mM sodium orthovanadate, 1 mM tetrasodium pyrophosphate, 2 mM PMSF, 10 μg/ml leupeptin, and 10 μg/ml aprotinin. Samples were then centrifuged at 16,000*g* for 10 min at 4°C, supernatants collected, and protein concentrations determined.

Equal amounts of protein samples were electrophoresed through a reducing sodium dodecyl sulfate polyacrylamide gel. Proteins in the gel were then electro-transferred to the Immobilon-FL Transfer Membrane (MilliporeSigma, Burlington, MA, USA). The membrane was blocked with Odyssey blocking buffer (LI-COR, Lincoln, NE, USA) for one hour at 25°C and incubated overnight with the rabbit anti-tau (C-terminal) antibody (MilliporeSigma) or the mouse anti-phospho-Tau (Thr181) Clone AT270 antibody (Thermo Fisher Scientific, Waltham, MA, USA) at 4°C. Washed membranes were then incubated with IRDye 680RD or IRDye 800CW (LI-COR) for one hour. Signals were obtained by using the Odyssey Infrared Imaging System (LI-COR).

### Immunofluorescence

Immunofluorescent analysis of tau protein and phosphorylated tau at threonine 181 was performed on 70% confluent human pulmonary artery smooth muscle cells. The cells were fixed with 4% paraformaldehyde for 15 minutes, permeabilized with 0.25% Triton X-100 for 10 minutes, and blocked with 5% BSA for 1 hour at room temperature. The cells were labeled with the rabbit anti-tau (C-terminal) antibody (MilliporeSigma) or the mouse anti-phospho-Tau (Thr181) Clone AT270 antibody (Thermo Fisher Scientific) at room temperature at dilution of 1:1000 in 1% BSA for 1 hour and then labeled with Alexa Fluor488 goat anti-mouse IgG secondary antibody at a dilution of 1:500 for 30 minutes at room temperature. F-actin was stained with Alexa Fluor 594 phalloidin. The slides were examined using the Olympus BX61 DSU Fluorescence microscope. Digital fluorescence micrographs were recorded and analyzed with the ImageProPlus software.

## Results

### Tau protein is expressed in the smooth muscle

Tau protein expressed in neuronal cells is well known to play key roles in neurological disorders. However, whether the smooth muscle expresses tau protein is not known. We found that tau protein is expressed in the smooth muscle layer of cerebral vessels of brain tissues from human patients who died of ischemic stroke as visualized by immunohistochemistry (Fig. 1A). Similar results showing the expression of tau protein in brain vascular smooth muscle were obtained in 3 patient samples. Tau protein was also detected by Western blotting in cell lysates prepared from cultured human brain vascular smooth muscle cells (Fig. 1B). Two different sequences of siRNA reduced the intensity of the band, confirming that this 40 kDa band is indeed tau protein expressed in smooth muscle cells (Fig. 1B). Similarly, tau protein expression was detected in human pulmonary artery smooth muscle cells (data not shown).

**Fig. 1:**
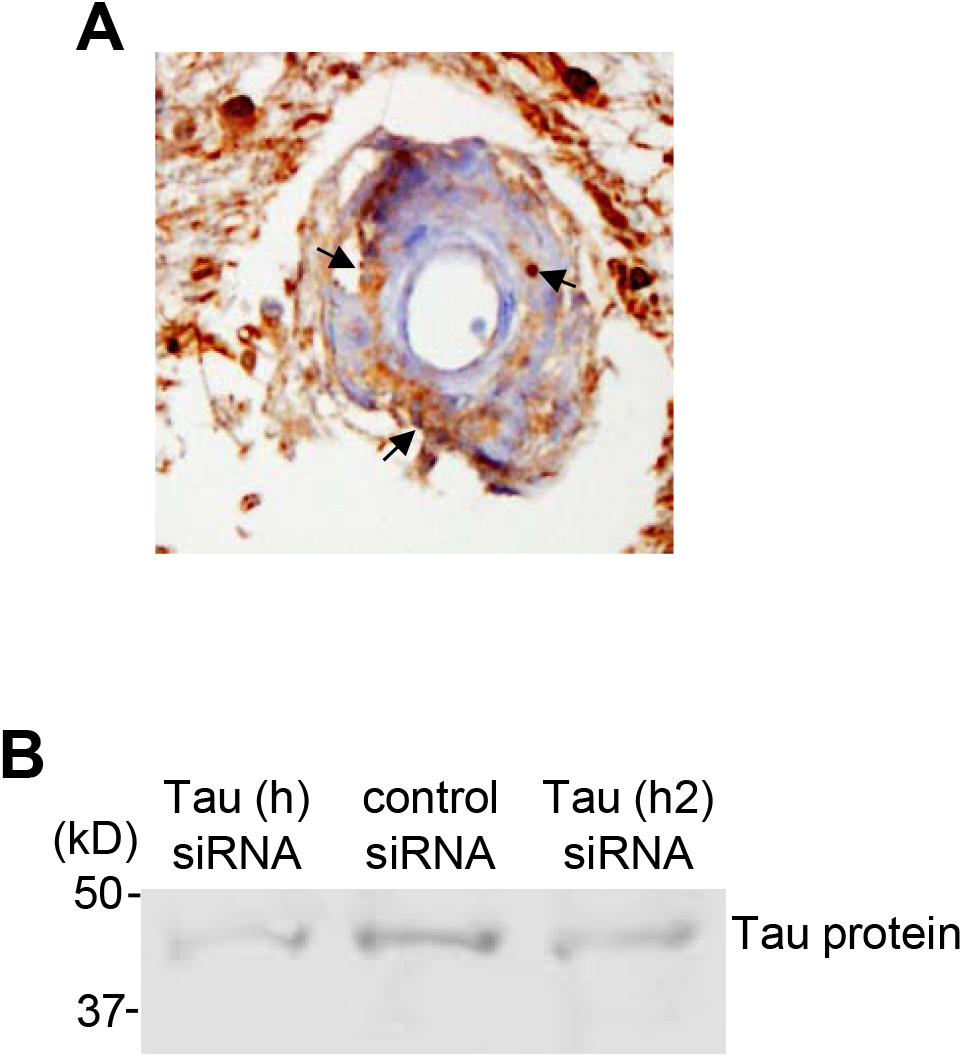
Tau protein is expressed in smooth muscle cells. (A) Immunohistochemistry shows that tau protein is expressed in the brain vascular smooth muscle tissues from patients with systemic hypertension died of ischemic stroke. (B) Cultured human brain vascular smooth muscle cells transfected with control, Tau (h) or Tau (h2) siRNA. 48 hours later, cell lysates were prepared. Western blotting with the Tau antibody.

### Tau protein is constitutively phosphorylated at threonine 181 in smooth muscle cells

We found that tau protein expressed in human brain vascular smooth muscle cells is constitutively phosphorylated at threonine 181 using Western blotting with a phospho-specific Tau antibody (Fig. 2). Similarly, the constitutively phosphorylated tau at threonine 181 was detected in human pulmonary artery smooth muscle cells of the lung vasculature as well as in human bronchial smooth muscle cells of the airways (Fig. 2).

**Fig. 2:**
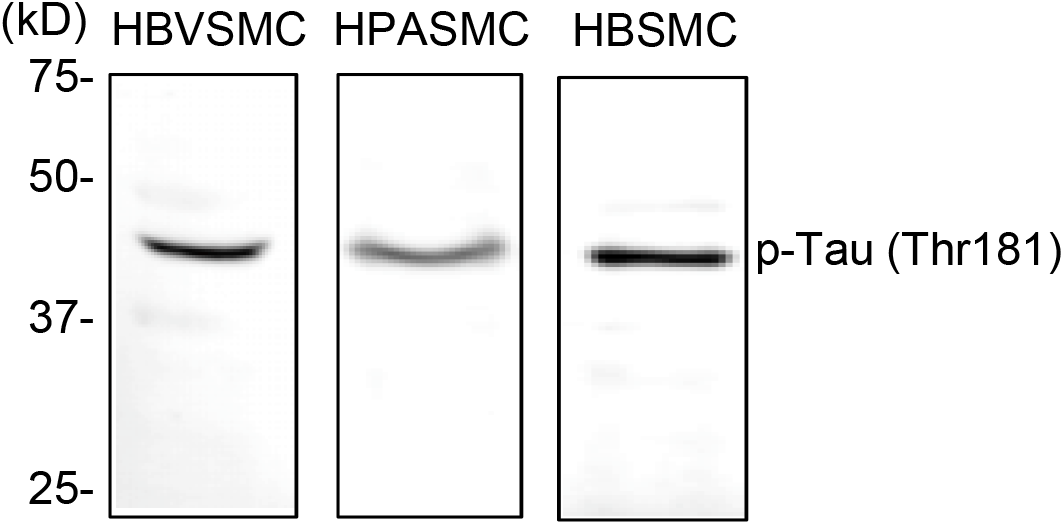
Threonine 181 of tau protein is constitutively phosphorylated in various smooth muscle cell types. Cell lysates prepared from cultured human brain vascular smooth muscle cells (HBVSMC), human pulmonary artery smooth muscle cells (HPASMC), and human bronchial smooth muscle cells (HBSMC) were subjected to Western blotting using the antibody against phosphorylated tau at Threonine 181.

### Cytoskeletal organization of tau and phosphorylated tau

Immunofluorescence using the tau antibody shows dispersed total tau protein expression in the cytoplasm of human pulmonary artery smooth muscle cells (Fig. 3A). By contrast, tau protein phosphorylated at threonine 181 was found to be associated with the microtubule and is well organized in the cytoplasm as determined by immunofluorescence staining using the phospho-specific tau antibody (Fig. 3B).

**Fig. 3:**
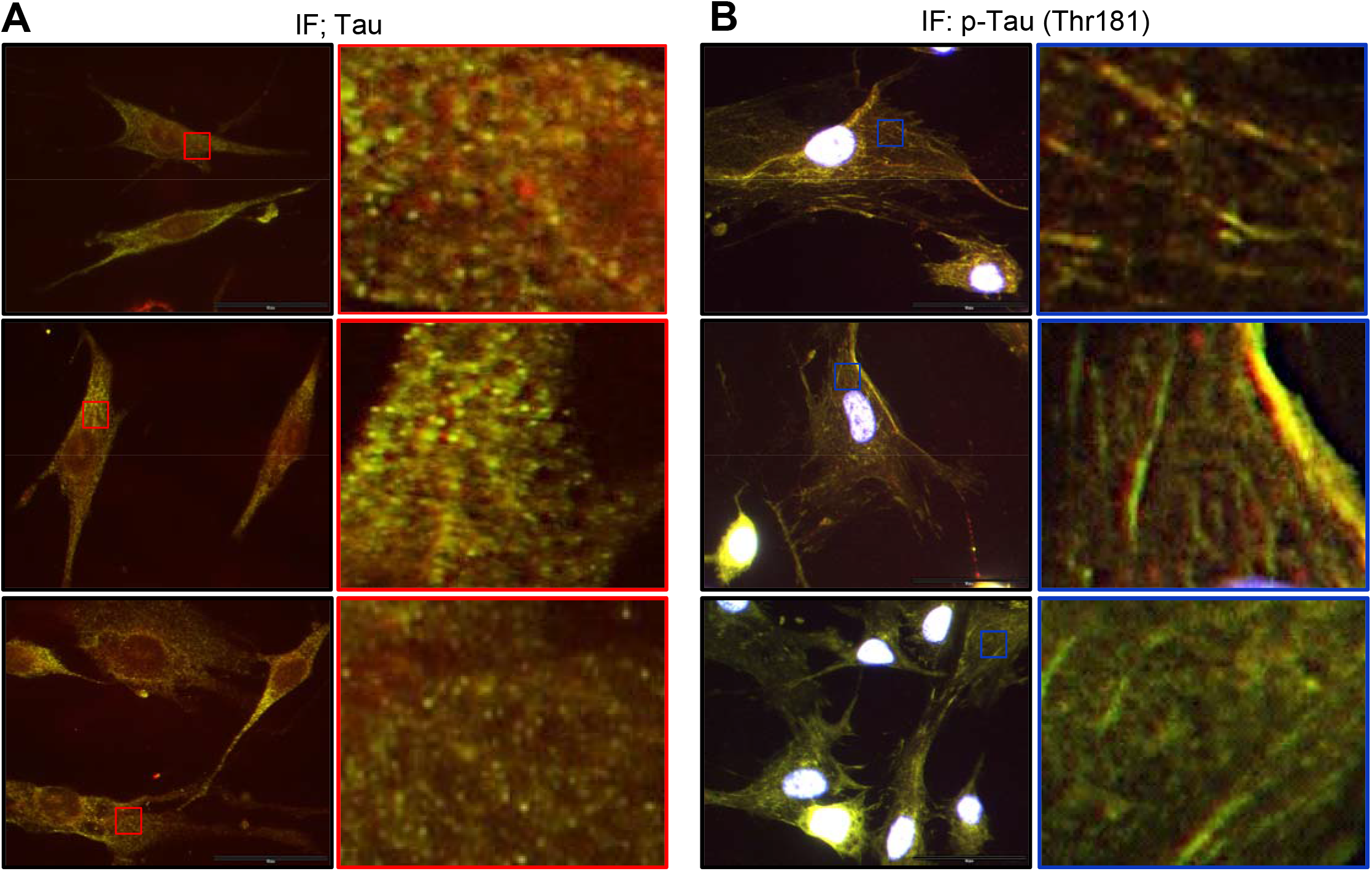
Tau protein that is phosphorylated at threonine 181 is highly organized in smooth muscle cells. Human pulmonary artery smooth muscle cells were subjected to immunofluorescence (IF) analysis using (A) the tau protein antibody and (B) phospho-specific tau (Thr 181) antibody (p-Tau). Areas indicated with squares are shown enlarged on the right side of each image. These images show a well assembled network of phosphorylated tau localized along the microtubules, while tau protein molecules that are not phosphorylated at threonine 181 are not well organized.

### Effects of calyculin A

Calyculin A is an inhibitor of protein phosphatase type 1 and 2A that should promote protein phosphorylation (Lerea et al., 2007; Nishikawa et al., 1994). Our Western blotting experiments revealed that calyculin A produced multiple higher molecular weight species that can be detected by the phospho-tau (Thr181) antibody in human pulmonary artery smooth muscle cells (Fig. 4A). These higher molecular weight species largely occurred at about 80 and 160 kDa and can be induced as early as 10 min of cell treatment with calyculin A and its formation continues to increase at 30 min (Fig. 4A). By contrast, subjecting cells to oxidative stress by the hydrogen peroxide treatment did not induce such higher molecular weight species (Fig. 4A). Molecular weights of 80 and 160 kDa are consistent with the multimers of tau protein. These may be the product of phosphorylation-dependent tau aggregation.

**Fig. 4:**
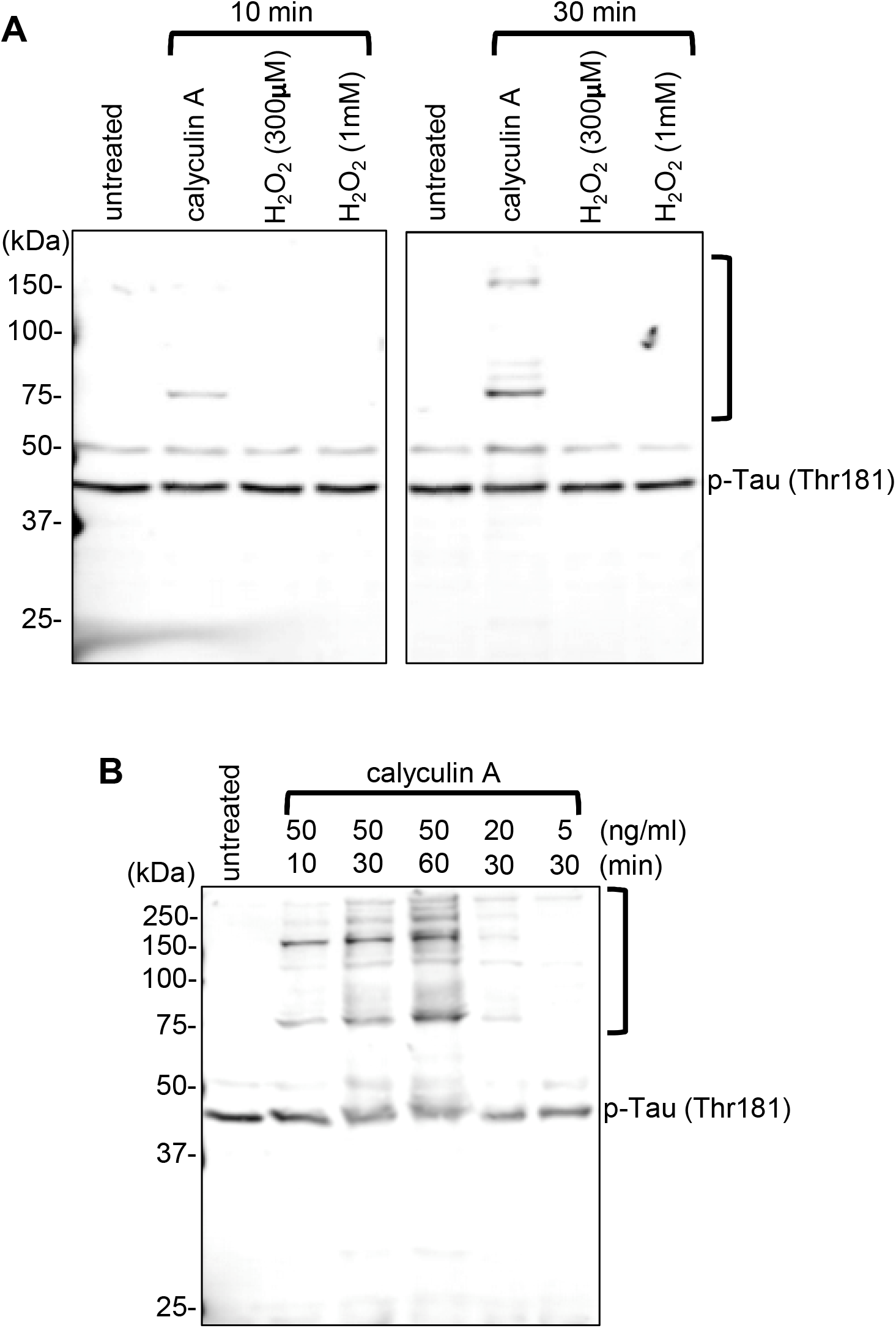
Calyculin A forms high molecular phosphorylated tau species. (A) Human pulmonary artery smooth muscle cells were treated with calyculin A (50 ng/ml) or hydrogen peroxide (H_2_O_2_) for durations indicated. (B) Human bronchial smooth muscle cells were treated with various doses of calyculin A for durations indicated. Cell lysates were subjected to Western blotting with the antibody against phosphorylated tau protein at threonine 181, p-Tau (Thr181).

Similarly, these high molecular weight species were also detected in airway smooth muscle cells treated with calyculin A. Fig. 4B shows that the treatment of human bronchial smooth muscle cells with calyculin A at 50 ng/ml caused for formation of higher molecular weight species of tau at 80, 120, 160, and perhaps 200 or 240 kDa in a time-dependent manner. The formation of high MW species occurred at as early as 10 min and continued to increase (Fig. 4B). Dose-dependence of calyculin A at 5, 20 and 50 ng/ml was also observed (Fig. 4B). We repeatedly and consistently observed the calyculin A-induced high molecular weight species formation as visualized using the phospho-tau (Thr181) antibody in smooth muscle cells in at least 10 experiments.

Immunofluorescence staining demonstrated that the treatment of human pulmonary artery smooth muscle cells with calyculin A disrupted the well organized cytoskeletal structure of tau phosphorylated at threonine 181 as seen in Fig. 3B and formed some protein aggregates as indicated by arrows (Fig. 5).

**Fig. 5:**
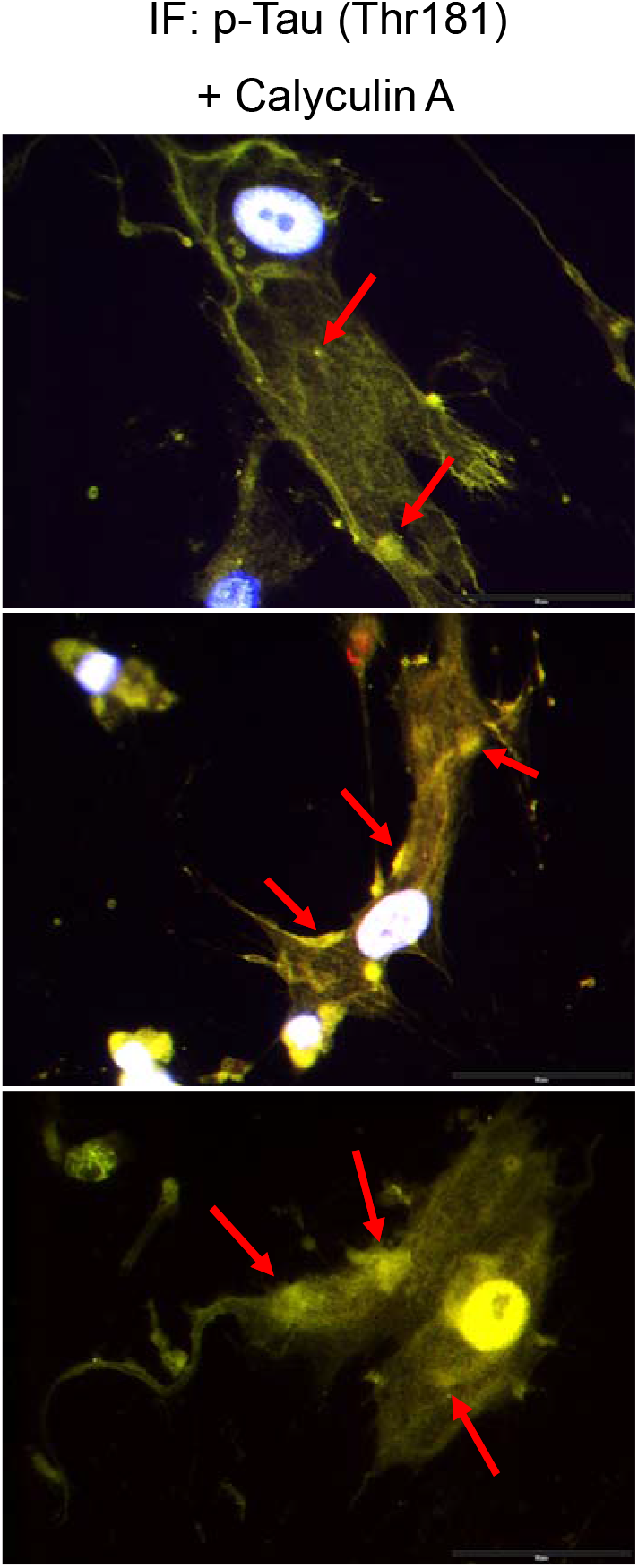
Calyculin A disturbs phosphorylated tau organization. Human pulmonary artery smooth muscle cells were treated with calyculin A (50 ng/ml) for 30 min and subjected to immunofluorescence (IF) analysis using the antibody against phosphorylated tau protein at threonine 181, p-Tau (Thr181). The representative images show the absence of well assembled microtubules as well as dispersed phosphorylated tau. The aggregation of phosphorylated tau is also visible as indicated with arrows.

## Discussion

Tau is a microtubule-associated protein that promotes the polymerization and assembly of microtubules and is considered to be one of the most important proteins in pathology of the central nervous system (Weingarten et al., 1975). It is located in the cellular compartment as well as in the interstitial fluid (Sotiropoulos et al., 2017). The abnormal accumulation of misprocessed tau is associated with various neurodegenerative diseases (Spillantini & Goedert, 2013). It has recently been shown that that tau has multiple functions in addition to the axonal microtubule assembly. Tau binds to nucleic acid and modulates gene expression and RNA stability (Bou Samra et al., 2017). In pathological conditions, tau causes DNA and RNA damage, nuclear disorganization, RNA and ribosome instability, and changes in protein expression (Tsartsalis et al., 2018; Meier et al., 2016; Frost et al., 2014). Tau may also modulate and impair cell signaling, contributing to altered receptors activities and cell death (Hamdane et al., 2005; Burnouf et al., 2013; Ittner et al., 2010).

Our finding in the present study showing that tau protein is expressed in various smooth muscle tissue/cell types opens up the possibilities that this protein may play pathophysiological roles in vascular, airway and gastrointestinal systems by exhibiting various mechanisms that this protein can elicit as described above. Like in nervous system, smooth muscle tau can be phosphorylated. This study specifically examined tau phosphorylation at threonine 181 in the smooth muscle, however, enormous amounts of investigations are now warranted to understand the role of various phosphorylation sites within the tau protein molecule.

It is noteworthy that phosphorylated tau at threonine 181 specifically assemble in a well organized fashion in smooth muscle cells, while most of tau protein molecules seem to be not so organized. These results highlight the potential importance of the threonine 181 phosphorylation of tau in smooth muscle biology.

Our finding that calyculin A promotes aggregation of tau is also noteworthy in that the tau aggregation that is seen in neurodegenerative diseases can also occur in smooth muscle cells. Further, our results consistently producing the higher molecular weight species corresponding to multimers of tau protein by calyculin A treatment of cells suggest the usefulness of this protein phosphatase inhibitor for the research of tau aggregation in brain cells in order to combat Alzheimer’s disease. It should be noted that Boban et al. (2019) reported that okadaic acid, another inhibitor of protein phosphatase type 1 and 2A, promoted the formation of high molecular weight tau species in SH-SY5Y cells. However, this tau species had a molecular weight of around 100 kDa that is not consistent with the product of tau aggregation.

In summary, the present study showed, for the first time, that smooth muscle cells express tau protein and that smooth muscle tau is capable of being phosphorylated and aggregated. Future studies further understanding smooth muscle tau protein may shed a light on normal cell biology as well as developing therapeutic strategies to combat a wide variety of diseases that affect smooth muscle cells.

## Acknowledgement

This work was supported in part by NIH (R01HL072844, R21AI142649, R03AG059554, and R03AA026516) to YJS. The content is solely the responsibility of the authors and does not necessarily represent the official views of the NIH.

